# OligoFlow: rapid and sensitive virus quantification using flow cytometry and oligonucleotide hybridization

**DOI:** 10.1101/2022.06.23.497420

**Authors:** James S. Paterson, Lisa M. Dann, Jessica Carlson-Jones, Sarah Giles, Connor McIvor, Peter G. Speck, James G. Mitchell

## Abstract

Flow cytometry is an established method for the detection and enumeration of viruses. However, the technique is unable to target specific viral species. Here, we present OligoFlow, a novel method for the rapid detection and enumeration of viruses by incorporating flow cytometry with species specific oligonucleotide hybridization. Using Ostried herpesvirus and dengue virus as model organisms, we demonstrate high-level detection and specificity. Our results represent a significant advancement in viral flow cytometry, opening the possibilities for the rapid identification of viruses in time critical settings.

## Main Body

Viruses are a fundamental element of the world in which we live. Their significant roles range from moderating ocean biogeochemical cycles to causing global pandemics, such as the SARS-CoV-2 outbreak^1,2^. The enumeration of viruses began with plaque assays^3,4^, a method that is still used. Soon after, viruses were enumerated by electron and epifluorescence microscopy^5,6,7,8^, providing more detailed information on the structure and shape of viruses and more accurate estimates of abundance than plaque assays. The first flow cytometry-based viral abundance measurements detected and discriminated between two types of viruses based on differences in their light scattering^9^. However, the introduction of new nucleic acid staining dyes in the late 1990’s led to the improvement in cytometric detection limits and has since transformed viral enumeration^10,11^, becoming a rapid and cost-effective method that continues to evolve^12^. In contrast to previous viral detection methods, flow cytometry routinely, rapidly and inexpensively counts single viruses from almost any sample type^13^. The technique characteristically takes tens of minutes for preparation and only minutes for a quantitative measurement. Current flow cytometry techniques enumerate viruses using generic nucleic acid fluorescent stains that target all viruses within a sample^14^. It is impossible to determine the presence or abundance of one specific viral species using current methods. More recently, the use of Fluorescence *in Situ* Hybridisation (FISH) combined with flow cytometry (Flow-FISH) for the identification and quantification of bacterial populations has become a commonly used method^15,16,17,18^. However, the use of Flow-FISH for viral species identification is yet to occur. Here, we develop OligoFlow for the rapid detection and enumeration of individual virus species (Fig. 1a) by incorporating methodological elements of Flow-FISH, without the need for long incubations, washing and concentrating steps. Two viral species, Ostreid herpesvirus 1 (OsHV-1) and Dengue virus (DENV), were used as model organisms for the development of OligoFlow.

**Fig. 1:**
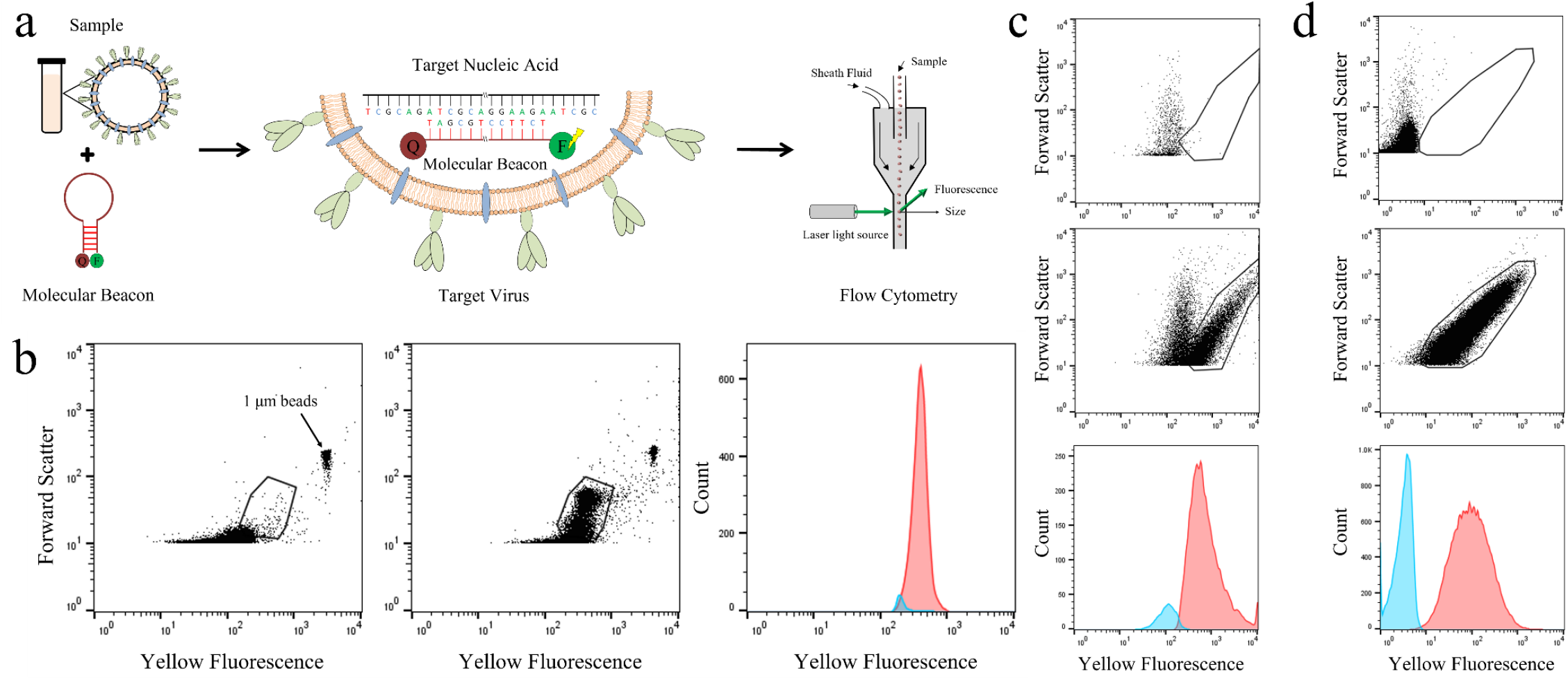
Principal method of OligoFlow and successful application with DENV and OsHV-1. **a**, Schematic overview of the OligoFlow methodology incorporating flow cytometry and oligonucleotide hybridization. Sequence specific Molecular Beacons fluorophores were designed to target the viral species of interest and then added to each sample. Samples were then incubated for fluorophore permeation through the viral capsid, then annealed for fluorophore binding to the complementary sequence. Analysis of each sample was carried out on the Merck Muse flow cytometer for the detection of yellow fluorescence, indicating successful probe binding, and forward scatter, a proxy for particle size. **b**, Initial analysis of pure DENV cultured samples on the Muse cytometer using generic nucleic acid fluorescent staining. The left cytogram depicts a “noise” sample with the addition of 1 μm fluorescent beads, while the cytogram on the right represents a DENV sample where positive events are detected in the gated region. Each black dot on all cytograms represents a detected particle. **c**, Detection of DENV using OligoFlow. The top cytogram represents a “noise” sample with minimal events within the gated region. The middle cytogram shows strong detection of DENV within the gated region, while the bottom histogram highlights the clear separation of yellow fluorescence signal between the “noise” and positive sample cytograms. **d**, Successful detection of OsHV-1 using OligoFlow, where the top cytogram represents a “noise” sample with no events present in the gated region. The middle cytogram shows an intense signal of OsHV-1 present in the gated region and the histogram shows separation of signal between the two cytograms.

The detection and enumeration of DENV was initially carried out using the nucleic acid stain SYTO Orange 81 (Molecular Probes) to evaluate the measured size and fluorescence sensitivity of the Muse cytometer (Merck). The Muse cytometer, an entry level machine equipped with a green laser (532 nm excitation), was chosen for the development of OligoFlow due to its ease of use and future potential applications. DENV was successfully detected with a clear cytometric signature using yellow fluorescence (576nm emission) and forward scatter detectors, where viral positive samples were discriminated against unstained control samples (Fig. 1b) to ensure true representation of viral detection. Concentrations calculated from within the gated signatures equated to 2.44 ± 0.37 × 10^6^ per mL^-1^. The supplied titre of DENV stock was approximately 10^6^ PFU per mL^-1^, highlighting the ability of the Muse cytometer for its viral detection capability and enumeration accuracy.

After successful detection and enumeration of DENV using a standard nucleic acid fluorescent dye, specific oligonucleotide fluorophores (Molecular Probes) for DENV and OsHV-1 were incorporated to combine flow cytometry with oligonucleotide hybridization. By combining species specific fluorescent probes and flow cytometry with short hybridization and preparation time, viral species of interest can be detected and enumerated rapidly and inexpensively regardless of sample type. We based our OligoFlow method on the denaturation and annealing aspects of PCR for successful probe attachment (Fig. 1a). The detection and enumeration of purified DENV and OsHV-1 samples was achieved using the OligoFlow method (Fig. 1c,d). The concentration of DENV using this method was 1.03 ± 0.45 × 10^6^ per mL^-1^, while OsHV-1 concentration was 2.89 ± 0.50 × 10^7^ per mL^-1^. The calculated concentrations of DENV using OligoFlow are consistent to concentrations identified with the same sample stained with SYTO Orange, demonstrating the accuracy and specificity of the OligoFlow method.

To test the applicability of OligoFlow on different sample types, Pacific oyster, *Crassostrea gigas*, tissue infected with OsHV-1 was extracted. Extracted OsHV-1 from tissue samples were successfully detected and enumerated using OligoFlow (Fig. 2a). The concentration of OsHV-1 from extracted oyster tissue varied greatly between individual oysters and ranged from below detection of 1 virus per mg^-1^ up to 1.29 × 10^6^ viruses per mg^-1^ of oyster tissue (Fig. 2a). These results indicate that OligoFlow is sensitive and specific enough to quantitatively determine the level of infection within an oyster population. The ability to integrate OligoFlow into time critical microbiological settings, such as cystic fibrosis exacerbations^19^, could provide rapid level of infection information to assist with alternative treatments and prevent death.

**Fig. 2:**
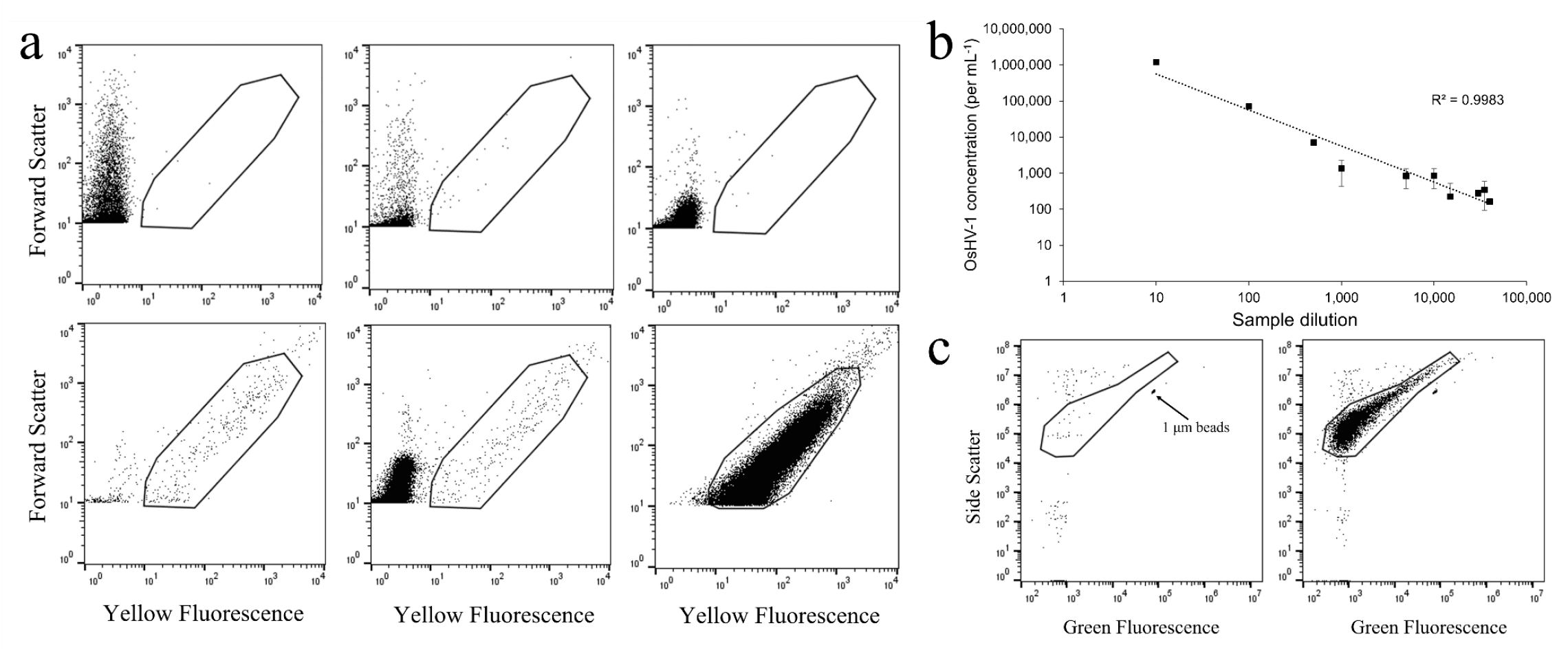
Specificity and sensitivity of OligoFlow with OsHV-1 extracted from oyster tissue. **a**, The detection of OsHV-1 in infected Pacific oysters, *Crassostrea gigas*, exhibited varying levels of viral load. The top left cytogram depicts a “noise” sample control, while all other cytograms represent individual oyster tissue extracts. The top right cytogram shows an oyster with little to no infection, while the bottom right cytogram shows an oyster heavily infected with OsHV-1. **b**, Serial dilution of extracted OsHV-1 from oyster tissue using OligoFlow exhibits a significant power trend across 10 dilutions ranging from 1:10 to 1:40,000. Dilution factors have not been factored in to calculate OsHV-1 concentration. The data points and error bars are means and standard errors, respectively, and the trend line is a power law. **c**, Analysis of extracted OsHV-1 on the Beckman Coulter CytoFLEX S cytometer. The left cytogram represents a “noise” sample with the addition of 1 μm fluorescent beads, while the right cytogram shows clear detection of OsHV-1 in the gated region, demonstrating the capability of OligoFlow across multiple flow cytometry platforms.

Next, to test the accuracy of OligoFlow we carried out a sample dilution experiment. Serial dilutions starting at 1:10 and up to 1:40,000 of OsHV-1 extracted from oyster tissue were prepared and analysed on the Muse cytometer. The concentration of OsHV-1 at 1:10 dilution was 1.20 ± 0.04 × 10^6^ per mg^-1^ (*n* = 3, ± SE) and at 1:40,000 dilution was 1.64 × 10^2^ per mg^-1^ (Fig. 2b). A power trendline was applied to all dilution samples and resulted in a R^2^ value of 0.9983 (p < 0.0001; Fig. 2b). This highlights a strong significant relationship between sample concentration and the accuracy of detection, indicating that the concentration of a virus has no effect on the accuracy of OligoFlow. The detection limit of our method is 14 viruses per mL^-1^, which is the detection of one virus particle from the analysis of a neat sample. This compares to qPCR methods having a reliable limit of detection as low as 384 copies of a target virus per mL^-1^ in a single reaction^20,21^.

Finally, the OligoFlow methodology was validated to ensure replicable and accurate use across multiple flow cytometer platforms. Purified OsHV-1 was analysed on a CytoFlex S cytometer (Beckman Coulter) equipped with violet (408nm), blue (488nm) and red (638nm) laser excitation and green (520nm) and orange (585nm) emission detectors. Samples were prepared identically with the samples analysed using the Muse cytometer. Successful detection and enumeration of OsHV-1 was achieved on the CytoFlex S (Fig. 2c) with a calculated concentration of 2.47 × 10^7^ per mL^-1^. This highlights that the OligoFlow method is not restricted to just one cytometer and can be transferred to any cytometry platform available without the loss of accuracy and detection capability.

The development of OligoFlow for the detection and enumeration of OsHV-1 and DENV has shown high accuracy, sensitivity and specificity with rapid and simple methodology that builds on the previous methods of bacterial Flow-FISH^17,18^. We have demonstrated that this method is also unrestricted by cytometer platform, making OligoFlow accessible to all flow cytometry users. The ability to identify and count a species of interest makes OligoFlow a method that could be utilized across a range of research, clinical and quality control settings. These features open the possibilities for the rapid identification of viruses in time critical settings.

## Methods

### Source of DENV and OsHV-1 virus

Purified culture samples of DENV were provided by the Virus Research Laboratory (Flinders University, Australia) at a titre of approximately 10^6^ PFU per mL^-1^. Initial samples of OsHV-1 were obtained from the Animal Health Laboratory, Department of Primary Industries, Parks, Water and Environment, Tasmania, Australia at a title of approximately 10^7^ PFU per mL^-1^. These samples were from extracted Pacific oyster tissue that was purified and suspended in filtered seawater. Samples were then snap frozen in liquid nitrogen and transported then stored at -80°C. Further samples of OsHV-1 were obtained from the South Australian Research Development Institute (SARDI) from an infection study where infected oyster tissue samples were collected. Tissue samples at an approximate size of 2 mm by 3 mm were cut from the oyster then placed in a microfuge tube with 500 µl of filtered and sterilised TE Buffer (10 mM Tris, 1 mM EDTA). The sample was then mechanically masticated for 5 min then centrifuged for 1 min at 6000 rpm. The supernatant containing OsHV-1 was separated and filtered through a 0.22 µm syringe filter and collected into a sterilized microfuge tube, snap frozen in liquid nitrogen and stored at -80°C until analysis.

### Development of a virus specific fluorophore

Fluorescent probes specifically for DENV and OsHV-1 were constructed through an online platform with Sigma-Aldrich Australia. Specifically, Molecular Beacons were developed for each virus that allowed sequence specific and highly sensitive detection. For OsHV-1, the primers C2F (CTCTTTACCATGAAGATACCCACC) and C6R (GTGCACGGCTTACCATTTTT) were used^22^. For DENV, the primers DENV5.1F (GCAGATCTCTGATGAATAACCAAC) and DENV3.2R (TTGTCAGCTGTTGTACAGTCG) were used^23^. All Molecular Beacons probes were attached with a HEX fluorophore (535nm excitation, 556nm emission) and a BHQ-1 quencher. The BHQ-1 molecule was used to quench the fluorescence of the HEX fluorophore until the attachment of the primer sequence occurs in the reaction.

### Detection and enumeration of DENV using SYTO Orange

Initial detection of DENV was carried out using SYTO Orange 81 (Molecular Beacons). This nucleic acid fluorescent stain was preferred due to the excitation (530 nm) and emission (544 nm) properties aligning closely to the Muse cytometer excitation and emission values. Samples for DENV were diluted 1:10 by adding 50 µl of virus sample to 450 µl of 0.02 µm filtered TE buffer (10 mM Tris, 1 mM EDTA). For each sample, 20 µl of SYTO Orange 81 (2.5 μM final concentration) was added then samples were incubated in the dark at 80°C for 10 minutes^14^.

### Sample preparation for viral species detection by OligoFlow

Samples for DENV and OsHV-1 were diluted 1:10 to assist in detection and not over saturate the cytometer detectors. Specifically, 25 µl of each virus was diluted in 225 µl of 0.02 µm filtered TE buffer (10 mM Tris, 1 mM EDTA). For each sample, 0.5 µl of each forward and reverse primer probe (20 nM final concentration) was added then samples were placed on a heat block at 80°C for 10 minutes then at 60°C for 5 minutes. Samples were then removed from the heat block at stored in the dark at room temperature until analysis.

### Flow cytometric analysis of DENV and OsHV-1

Samples of DENV and OsHV-1 were analysed on a Muse cytometer (Merck). Prior to each session calibration beads were prepared and run in triplicate for quality control and calibration of the volumetric sensor. After the preparation and incubation of each sample as described above, individual samples were loaded into the machine and analysed for 2 minutes or 50,000 events. After the analysis of each sample the cytometer was rinsed with sterile MilliQ water to eliminate any sample crossover. Further samples of OsHV-1 were analysed on the CytoFlex S cytometer (Beckman Coulter). Calibration beads were run before each session and samples were analysed for 2 minutes.

### Serial dilution of OsHV-1 for analysis by OligoFlow

Samples of OsHV-1 extracted from infected oyster tissue were prepared in a serial dilution, whereby OsHV-1 was diluted in 0.02 μm filtered TE Buffer (10 mM Tris, 1 mM EDTA) at dilutions of 1:10, 1:100, 1:500, 1:1,000, 1:5,000, 1:10,000, 1:15,000, 1:30,000, 1:35,000 and 1:40,000. Each dilution was prepared and analysed in triplicate using the Muse cytometer.

## Data analysis

Raw flow cytometry (FCS) files were exported from the Muse and CytoFLEX S cytometers and analysed in FlowJo software (Becton Dickinson). Populations of DENV and OsHV-1 were discriminated based on differences in forward scatter (FSC) and yellow fluorescence on the Muse, and differences in side scatter (SSC) and green fluorescence on the CytoFLEX S. Concentrations were calculated using the raw FCS files combined with the calibrated analysed volume of each sample recorded by the Muse and CytoFLEX cytometers.

## Acknowledgements

This project was funded by the Fisheries Research and Development Corporation (FRDC) under project number 2016-806. We thank Graeme Knowles from the Animal Health Laboratory, Department of Primary Industries, Parks, Water and Environment, Tasmania and to Dr Marty Deveney from SARDI Aquatic Sciences for sourcing and providing oyster samples infected with OsHV-1 for research to be conducted on. We also thank Associate Professor Jill Carr from the College of Medicine and Public Health, Flinders University for providing Dengue virus samples to conduct experiments.

